# Characterizing a Low-Density Neutrophil gene signature in acute and chronic infections and its impact on disease severity

**DOI:** 10.1101/2024.11.01.621545

**Authors:** Matheus Aparecido de Toledo, João Victor Souza de Lima, Reinaldo Salomão, Giuseppe G. F. Leite

**Affiliations:** Division of Infectious Diseases, Department of Medicine, Escola Paulista de Medicina, Universidade Federal de São Paulo, São Paulo, Brazil

**Keywords:** Low-density neutrophils, transcriptomics, proteomics, sepsis, infection severity, biomarkers, gene expression, PBMCs

## Abstract

Low-density neutrophils (LDNs) or polymorphonuclear myeloid-derived suppressor cells (PMN-MDSC) are involved in the pathogenesis of cancer, autoimmune and infectious diseases. They are crucial in the host-response to invading pathogens, especially during acute illness, and are associated with poor prognosis in many infectious diseases. However, their gene expression profile and contribution to disease outcomes are not well described. We conducted a meta-analysis of gene expression datasets from peripheral blood mononuclear cells (PBMCs), focusing on patients with viral and bacterial infections. We identified a consensus set of 2,798 differentially expressed genes. Among these, 49 genes were commonly found in both the neutrophil degranulation pathway and the granule lumen-specific community. To validate this signature, we evaluated its expression in RNA-seq datasets, finding consistent upregulation of 24 genes in severe infections, 17 of them overlapped with genes overexpressed in CD16^int^ cells. We also investigated the abundance of LDN-related proteins in a PBMC proteomics dataset from a cohort of sepsis and septic shock patients, identifying 13 proteins with significantly higher levels in sepsis and septic shock patients compared to healthy controls. In conclusion, our study identified a pattern of 17 upregulated LDN genes, common to PBMC-transcriptome and RNA-seq, and up-regulated in CD16^int^, associated with acute infections and severe clinical outcomes, marking the first time these genes have been collectively presented as a potential signature of LDNs in relation to disease severity. Further research with prospective cohorts is needed to validate this LDN signature and explore its clinical implications.

**Summary Sentence:** Meta-analysis revealed a 17-gene LDN signature associated with severity in acute infections, providing potential biomarkers for clinical outcomes in infectious diseases.

## 1. INTRODUCTION

Neutrophils are the most frequent type of immune cells in human blood, playing a crucial role in the innate immune system [1]. They employ various strategies, including the production of reactive oxygen species (ROS), the release of antimicrobial proteins, and the regulation of the inflammatory response, to provide a first line of defense against infection. [1, 2]. Even though these are well known functions, it has been shown that there is a large heterogeneity in neutrophil function and life span and these areas are still being actively investigated [1–3].

Recently, researchers have shown growing interest in a subpopulation of neutrophils known as low-density neutrophils (LDNs) or polymorphonuclear myeloid-derived suppressor cells (PMN-MDSC) [4–9]. These cells are characterized in peripheral human blood samples by their co-fractionation with peripheral blood mononuclear cells (PBMCs) during density gradient centrifugation, resulting in their designation as “low-density” [10]. LDNs have been detected in a variety of infectious illnesses, including Ebola [11], COVID-19 [4–7], tuberculosis [12, 13], and sepsis [8, 9]. Elevated LDN-related genes, proteins, and cell counts are associated with poor prognosis [5, 14], reflecting a common feature of emergency granulopoiesis in response to diverse pathogenic conditions.

LDNs are heterogeneous, consisting of both immature and mature neutrophils with segmented, banded, and myelocyte-like nuclei [15, 16]. They exhibit immunosuppressive properties in cancer and infectious diseases, achieved through ROS generation and arginase-1 release, which inhibits T-cell proliferation [17–19]. In contrast, during autoimmune disease, LDNs act as a pro-inflammatory mediator, promoting spontaneous neutrophil extracellular trap (NET) formation by neutrophils, enhancing the production of pro-inflammatory cytokines, and contributing to endothelial toxicity [16, 20, 21].

The primary objective of this study was to assess the expression of LDN-related genes in infectious diseases and their potential impact on the outcome of patients. To achieve this, we conducted a comprehensive analysis of gene expression datasets from PBMCs of patients with acute and chronic infections. Our findings demonstrated upregulation of LDN-related genes during acute infection episodes. Based on these observations, we further investigated the expression of LDN-related genes in RNA-seq datasets and their corresponding protein levels in a cohort of septic patients to better understand their potential roles in the disease context.

## 2. MATERIAL AND METHODS

### 2.1 Microarray Datasets Selection and Data Analysis

The microarray datasets for this study were obtained from the Gene Expression Omnibus (GEO) database (https://www.ncbi.nlm.nih.gov/geo/) and manually curated based on the following inclusion criteria: i) datasets involving human peripheral blood mononuclear cells (PBMCs); ii) presence of infection-related information, including details about the etiological agents and patient status; and iii) inclusion of healthy control groups. Exclusion criteria included: i) non-human data and ii) datasets with fewer than five patients or fewer than three healthy controls.

Applying these criteria led to the identification of nine datasets for subsequent analysis. Among these, the GSE6269 dataset included four distinct subsets, each representing a different pathogen: *Staphylococcus aureus*, *Streptococcus pneumoniae*, *Escherichia coli*, and *Influenza A*. Similarly, the GSE34205 dataset comprised two subsets, one related to *Influenza* and the other to *Respiratory Syncytial Virus* (RSV). Each subset was analyzed independently, leading to a total of 13 analyses (see **Table 1** for details; additional information is available in **Supplementary Material 1: Table S1**). Some datasets included data from patients with active or latent infections, with or without treatment, at varying stages of treatment, and under different treatment regimens. To ensure consistency, we limited our analysis to data from patients with active infections who had not received any form of treatment.

**Table 1.** Description of the PBMC microarray datasets selected from GEO.

| Disease | Dataset ID | Patients (N) | Controls (N) | Acute or chronic |
| --- | --- | --- | --- | --- |
| Hepatitis B | GSE58208 | 12 | 5 | Chronic |
| TB | GSE54992 | 9 | 6 | Chronic |
| Malaria | GSE5418 | 15 | 22 | Acute |
| HIV | GSE2171 | 22 | 12 | Chronic |
| HIV | GSE77939 | 5 | 4 | Chronic |
| Influenza A | GSE6269 | 18 | 6 | Acute |
| E. coli | GSE6269 | 29 | 6 | Acute |
| S. aureus | GSE6269 | 31 | 6 | Acute |
| S. pneumoniae | GSE6269 | 13 | 6 | Acute |
| RSV | GSE34205 | 51 | 10 | Acute |
| Influenza | GSE34205 | 28 | 12 | Acute |
| Rotavirus | GSE2729 | 10 | 8 | Acute |
| HCV | GSE40184 | 10 | 6 | Chronic |

Abbreviations: TB: Active/Latent Tuberculosis, CHB: Chronic Hepatitis B, HIV: Human Immunodeficiency Virus, E. coli: Escherichia coli, S. pneumoniae: Staphylococcus aureus, S. pneumoniae: Streptococcus pneumoniae, RSV: respiratory syncytial virus and HCV: Hepatitis C Virus.

We analyzed each dataset using R statistical software (version 4.0.1), performing differential expression analysis with the LIMMA method. Gene symbols corresponding to each probe were obtained from the GEO annotation file. In cases where multiple probes represented the same gene, the probe with the highest mean expression level in each dataset was selected. For each comparison, we computed the log2 fold change (FC) and the corresponding confidence intervals. A meta-analysis was then performed using the MetaVolcanoR package [22], generating a consensus score based on a p-value cutoff of ≤ 0.05, with genes required to be identified in at least 5 of the 13 analyses. The HIV dataset (GSE77939) did not yield any genes with p-values ≤ 0.05 and was therefore excluded from further analysis.

Additionally, we applied the Random Effects Model (REM) using MetaVolcano to summarize gene log2FC across multiple studies, accounting for inter-study variance. The REM estimated a summary p-value (REM p-value), representing the likelihood that the REM log2FC significantly differs from zero [22].

### 2.2 Global Infection Protein-Protein Network, Multiscale Community Detection and Overrepresentation Pathway Analysis

The gene list generated from the MetaVolcanoR analysis using the vote-counting function was used to build a protein-protein interaction network (PPIN) in Cytoscape (version 3.10). The global infection PPIN was created based on interactions from the STRING database, using the Cytoscape stringApp (v. 1.5.1) for *Homo sapiens*, with a confidence cutoff score set to 0.7, and no additional interactors included. A multiscale community detection analysis was then performed to group the genes into communities using the CyCommunityDetection.app in Cytoscape. The parameters for this analysis were as follows: Algorithm: OSLOM, Weight Column: STRING score, Random Number Seed: 1, with all other parameters set to their default values. Functional enrichment of the identified communities was conducted using the “g: Profiler” algorithm. In addition, pathway Overrepresentation Analysis (ORA) was performed on the gene list using DAVID DB, with Reactome pathways as the background (FDR < 0.05).

### 2.3 Validation of Gene Expression Portrait of LDNs Using RNA-seq Transcriptomic

To validate the gene expression profile associated with LDNs and explore potential associations with disease severity, we selected seven RNA-seq transcriptomic datasets, primarily from cases of acute viral infections, using the same selection criteria as applied for the microarray datasets (**Table 2**; additional information is available in **Supplementary Material 1: Table S2**). Raw count matrices were obtained directly from GEO and imported into the R statistical environment. Genes with low expression levels were filtered out to improve robustness and reduce noise. Differences in gene expression between cases and controls were assessed using Benjamini–Hochberg (BH)-adjusted moderated t-statistics in DESeq2 [23]. The resulting gene list was subsequently imported to MetaVolcanoR, where we applied REM approach for meta-analysis.

**Table 2.** Description of the PBMC RNA-seq datasets selected from GEO.

| Disease | Dataset ID | Comparison (N) |
| --- | --- | --- |
| Sepsis | GSE216902 | Non-responder (14) vs. Responder (23) |
| COVID-19 and Sepsis | GSE199816 | Sepsis (18) vs. COVID-19 (9) |
| COVID-19 | GSE206263 | Severe (6) vs. Mild (5) |
| COVID-19 | GSE152418 | Severe (8) vs. Moderate (4) |
| COVID-19 | GSE157859 | Severe (5) vs. Mild (4) |
| Dengue | GSE178240 | With plasma leakage (55) vs. Without plasma leakage (33) |
| Dengue | GSE94892 | Dengue shock (15) vs. Dengue fever (07) |

Additionally, we downloaded and analyzed an RNA-seq dataset of CD16^hi^ and CD16^int^ LDNs from three patients with severe COVID-19, as well as normal-density neutrophils (NDNs) from healthy donors.

### 2.4 Sensitivity Analysis

A sensitivity analysis was conducted using the microarray data, this time focusing exclusively on acute patient data. By restricting the analysis to this subset, we were able to perform a more targeted REM MetaVolcanoR analysis, ensuring that the results were robust and accurately represented gene expression patterns in acute infection cases.

### 2.5 Validation of Portrait of LDNs Using Proteomics

We investigated the proteins related to LDNs from a previously published cohort of PBMC proteomics by our research group [8]. Using the previously normalized data, we analyzed the abundance of proteins encoded by the genes identified in section 2.3. For a detailed description of the proteomics methods, see Leite, Ferreira [8]. The dataset includes nine healthy controls and 24 sepsis patients, 11 of whom had septic shock at the time of sample collection. We used Welch’s t-test to evaluate protein abundance in healthy controls versus sepsis patients (n = 24). In addition, ANOVA followed by Tukey’s HSD test was performed to evaluate protein levels in healthy controls, sepsis patients, and septic shock patients.

## 3. RESULTS

### 3.1 LDN signature in viral and bacterial, acute and chronic infections using microarray datasets

We analyzed gene expression in 253 patients with different infections and 91 healthy controls across 13 datasets. The MetaVolcanoR R package was used to process the gene lists generated from these analyses. The gene list produced by the vote-counting function was filtered to include only genes found in at least five datasets, resulting in a final list of 2,798 consensus differentially expressed genes (**Figure 1A**, and **Supplementary Material 1: Table S3**). This list was subsequently subjected to ORA using the Reactome database, which generated 25 significantly overrepresented pathways. Notably, “neutrophil degranulation”, with 179 genes, was one of the top five most significant pathways (**Figure 1B**).

**Figure 1.**
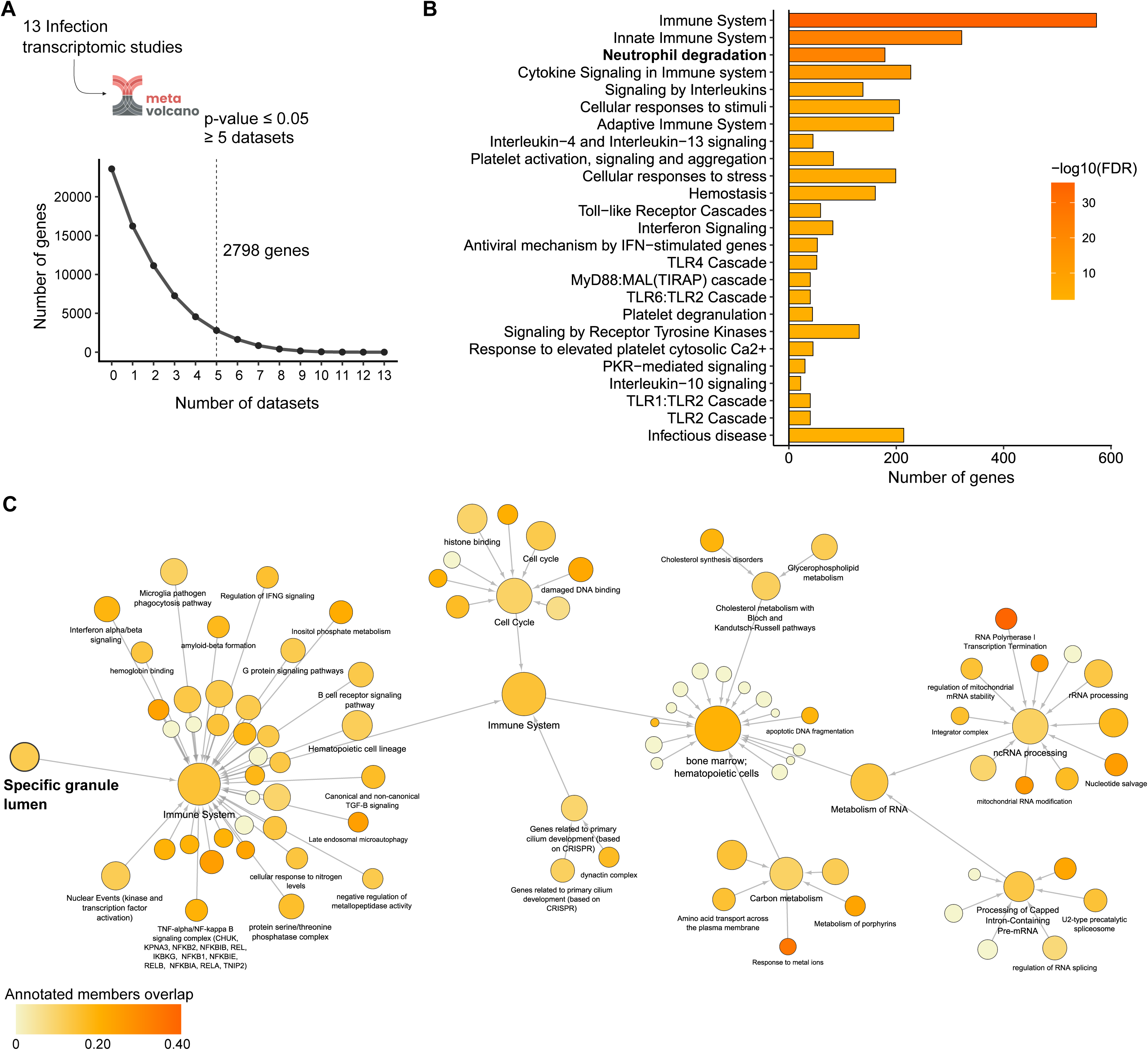
Meta-analysis of PBMC transcriptome studies in patients with infections. **Description Figure 1: A**) Number of differentially expressed genes across various infection conditions compared to healthy controls. The lines represent the number of genes identified by the vote-counting function in MetaVolcanoR (p-value < 0.05) present in one or more microarray datasets (x-axis). The number of genes found in at least five studies is indicated (2,798 consensus differentially expressed genes). **B**) Bar plot showing the overrepresentation analysis of Reactome pathways based on the consensus list of differentially expressed genes; and **C**) The multiscale community network of the global infection protein-protein interaction network (PPIN). Node size represents the number of genes annotated in each process, while the color scale indicates the ratio of overlapping genes in each annotation.

We then built a communities interaction network using the CyCommunityDetection app and explored the associated biological functions (**Figure 1C**). This network revealed 69 distinct communities, many of which were related to immune system processes and metabolism. Between these, we identified a module termed “specific granule lumen” consisting of 120 genes.

A Venn diagram comparing the genes of the neutrophil degranulation pathway with the granule lumen-specific community identified 49 genes common to both annotations (**Figure 2A; Supplementary Material 2: Table S4**). This set includes genes previously associated with LDNs in the literature, such as *AZU1, CAMP, CTSG, DEFA4, ELANE, MMP8, MPO, OLFM4, PLAUR, PTX3, RNASE2, RNASE3, S100A11,* and *S100A12* [5, 7, 8, 11, 14].

**Figure 2.**
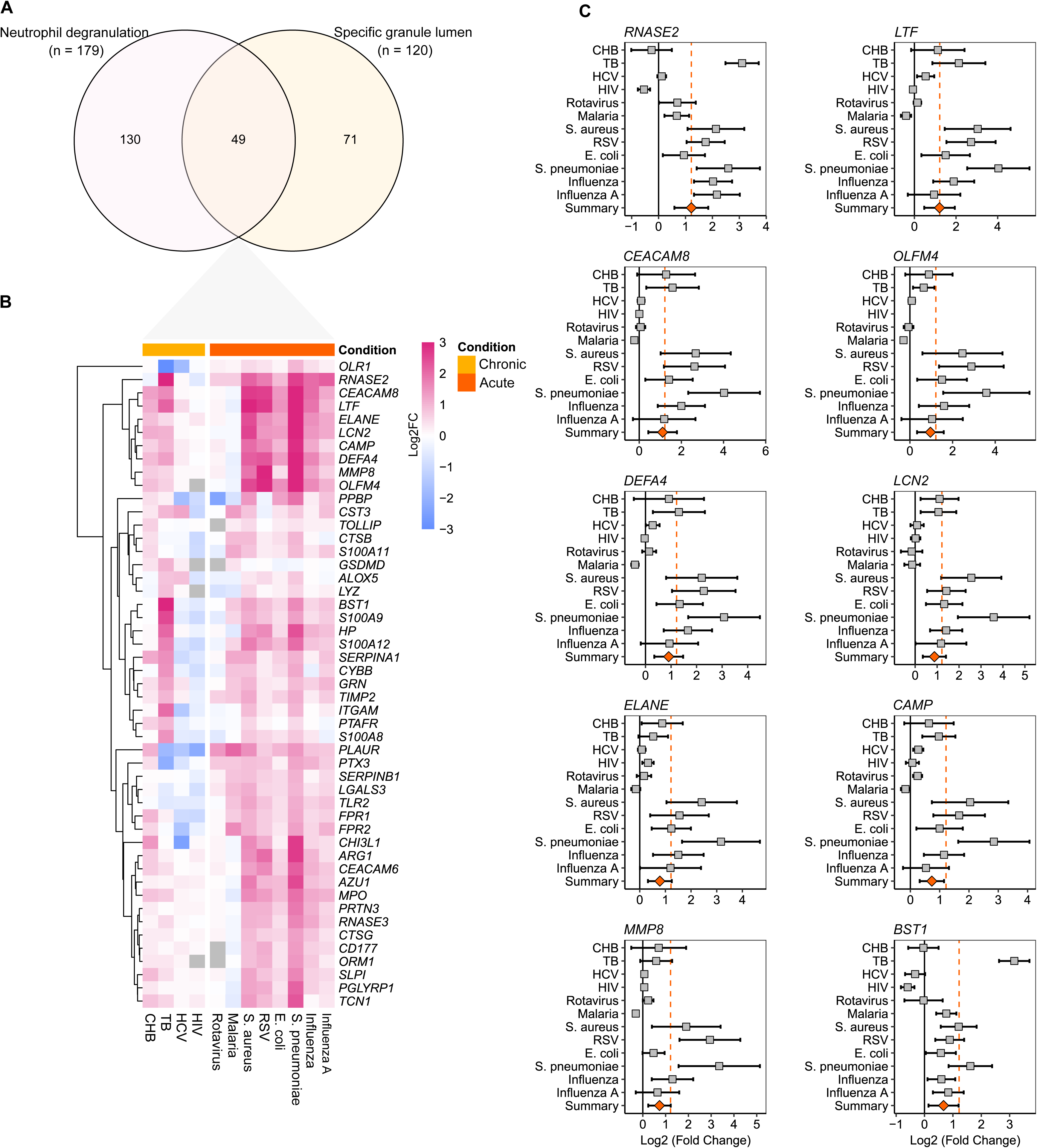
Low-density neutrophil (LDN)-related genes in various infections. **Description Figure 2:** Venn diagram showing the 49 consensus differentially expressed genes common to the two enrichment methods: “Neutrophil degranulation,” identified through overrepresentation analysis using the Reactome database, and “Specific granule lumen,” obtained from the interaction communities network using the CyCommunityDetection app. **B)** HeatMap showing variation in the magnitude of expression of these genes, Log2FC represents the variation in comparison between infected patients and healthy controls; and **C)** Forest plot displaying the top 10 genes with the highest REM log2FC and REM p-value ≤ 0.05, as determined by MetaVolcanoR.

Overall, the expression of these 49 genes was generally upregulated in patients with infections compared to healthy controls (**Figure 2B**). Among these, 31 genes showed a significant REM p-value ≤ 0.05 (**Supplementary Material 2: Table S5**). The top 10 REM log2FC with a REM p-value ≤ 0.05 are presented in **Figure 2C**. Most of these genes were upregulated in acute infections such as *S. aureus*, RSV, *E. coli*, *S. pneumoniae*, and influenza. In chronic infection states, the gene expression showed a positively regulated profile in CHB and TB, while mixed profile in HCV and HIV infections.

Collectively, these results suggest that LDN-related genes tend to be up-regulated, especially in acute infection states. To further explore this observation, we performed a sensitivity analysis using the REM MetaVolcano, focusing on datasets classified as acute. This analysis showed 43 out of the 49 genes with a REM p-value ≤ 0.05, all exhibiting a positive REM log2FC (**Supplementary Material 2: Table S6**).

### 3.2 Validation of Gene Expression Portrait of LDNs and Potential Associations with Severity

Previous studies have reported an increased presence of LDNs in severe infections [9, 24]. To assess if the 49 genes are upregulated in severe cases of different acute infections, we analyzed gene expression in seven RNA-seq datasets comprising 206 patients. Notably, we identified 24 genes with a REM p-value ≤ 0.05, and all exhibited a positive REM Log2FC (**Figure 3A**; **Supplementary Material 2: Table S7**).

**Figure 3.**
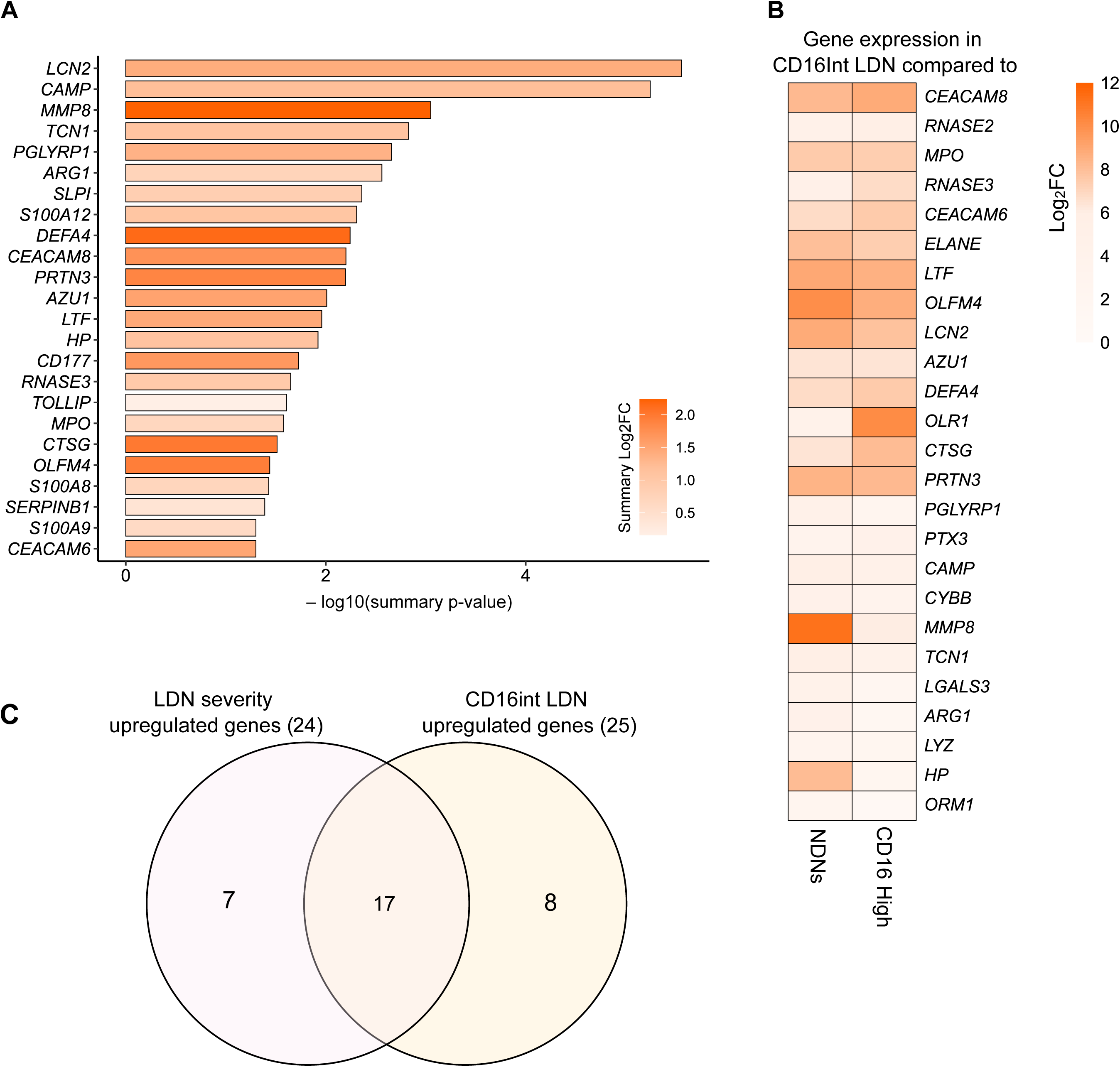
Meta-analysis of genes in RNA-seq datasets comparing patients with infections stratified by severity. **Description Figure 3: A)** Bar plot displaying the 24 genes with REM p-value ≤ 0.05, along with their respective REM log2FC (color scale). **B)** Heatmap showing the magnitude of individual genes differences in the CD16^Int^ LDNs relative to the CD16^hi^ LDNs and NDNs, expressed as log2FC; and **C)** Venn diagram showing the overlap between the results of the severity analysis and the CD16^Int^ LDNs analysis.

We also explore the presence of the 49 genes in an RNA-seq dataset of LDNs classified as CD16^hi^ and CD16^int^ from three patients with severe COVID-19, along with NDNs from healthy donors. The CD16^int^ LDN population has been classified as band-shaped nucleus, resembling immature neutrophil morphology [5]. We conducted two main comparisons: the first between CD16^int^ and CD16^hi^ LDN populations in COVID-19 patients, and the second between CD16^int^ LDNs and NDNs. In both comparisons, we found that 25 genes were significantly upregulated in CD16^int^ LDNs, with a Log2FC of 1.0 or higher and a BH-adjusted p-value of 0.05 or less. (**Figure 3B**). Among these, 17 genes (including *LCN2, CAMP, MMP8, TCN1, PGLYRP1, ARG1, DEFA4, CEACAM8, PRTN3, AZU1, LTF, HP, RNASE3, MPO, CTSG, OLFM4,* and *CEACAM6*) overlapped with the 24 genes previously identified as upregulated in LDNs associated with disease severity (**Figure 3C**).

These genes are intricately involved in key biological processes central to neutrophil functions. *CAMP, DEFA4, RNASE3, MPO, CTSG, AZU1, LTF, MMP8*, *PRTN3, OLFM4, LCN2 (NGAL)* and *PGLYRP1*, are involved in antimicrobial activity through peptides and enzymes stored in granules, participate in neutrophil degranulation, and facilitate pathogen clearance [6, 25–27]. *LCN2*, *LTF*, and *HP* limit bacterial growth by sequestering iron, which is essential for bacterial proliferation, and contribute to antimicrobial defense [27]. *CEACAM6* and *CEACAM8* facilitate cell adhesion and activation, supporting neutrophil migration during inflammation [6, 27]. *TCN1* is a vitamin B12-binding protein which regulates cobalamin homeostasis [27], while *ARG1* and *OLFM4* plays a role in immune regulation and modulation [27–29]. Together, these genes play a crucial role in various neutrophil functions.

### 3.3 Abundance of LDN-Related Proteins in Patients with Sepsis and Septic Shock

The PBMC proteomics cohort included 24 patients (13 with sepsis and 11 with septic shock) and 9 healthy controls. Out of the 17 genes analyzed, 13 corresponding proteins were identified. First, we directly compared the 24 patients with sepsis to the 9 healthy controls; 10 of these 13 proteins demonstrated significantly higher abundance in patients with sepsis compared to healthy controls (**Supplementary Material 2: Table S8**). These changes in abundance of these 10 proteins were also statistically significant when comparing the three groups (sepsis, septic shock, and healthy controls). Most proteins were higher in abundance in both sepsis and septic shock patients compared to healthy controls (**Figure 4**). Notably, proteins such as lactotransferrin (LTF), azurocidin (AZU1), and cathelicidin antimicrobial peptide (CAMP) were particularly elevated in sepsis patients, with CAMP being the only protein exhibiting differential abundance between sepsis and septic shock. Thus, further investigation with more patients is required for determining an association between LDN-related proteins and the severity of infectious diseases.

**Figure 4.**
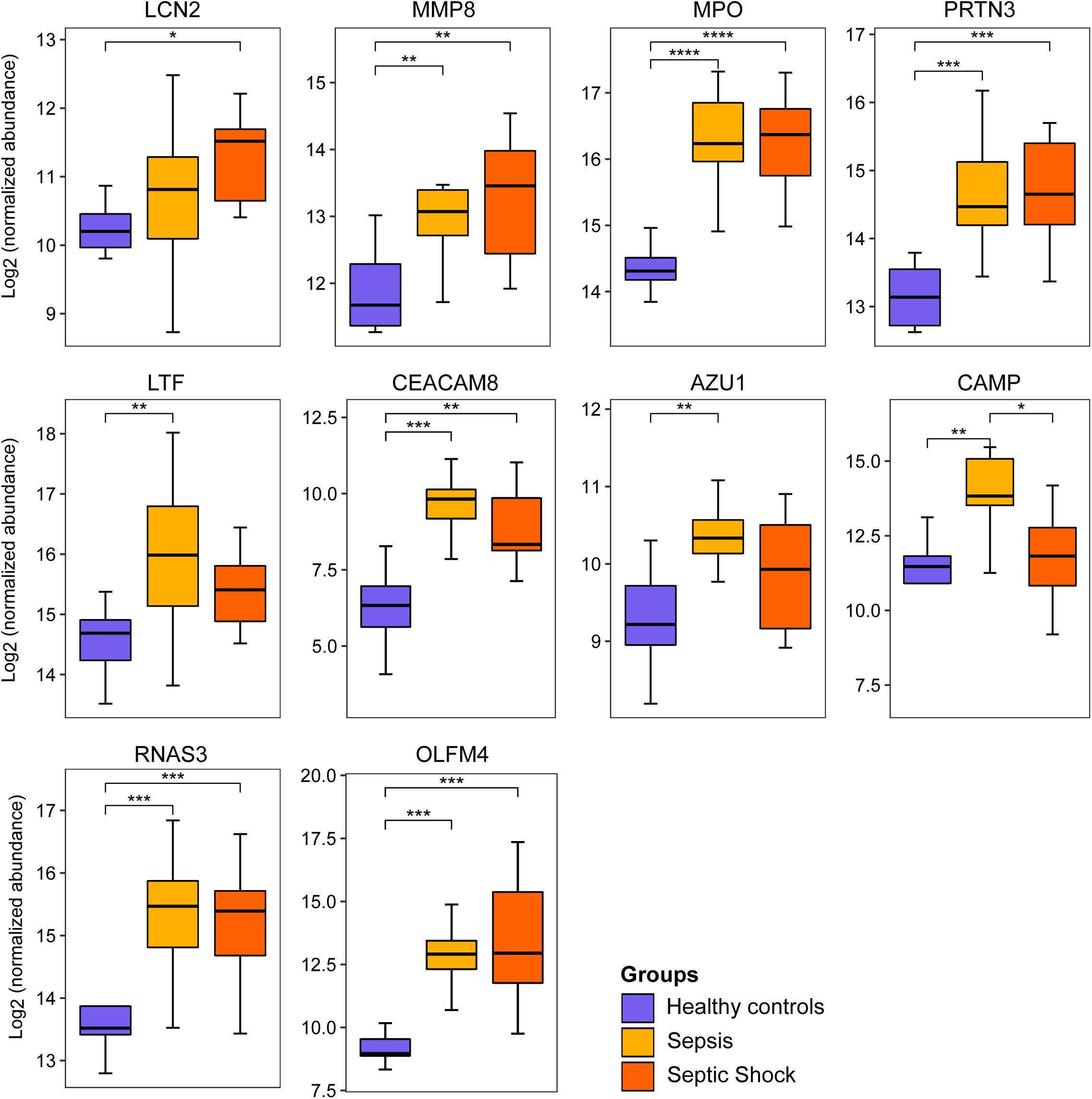
Abundance of LDN-Related proteins in sepsis and septic shock patients compared to healthy controls. **Description Figure 4 :** Boxplots depicting the abundance levels of 10 LDN-related proteins in patients with sepsis, septic shock, and healthy controls. Statistical analyses were performed using ANOVA followed by Tukey’s HSD test, with correction by the Benjamini– Hochberg method. Significance levels are indicated as follows: **** p < 0.0001, *** p < 0.001, ** p < 0.01, * p < 0.05.

## 4. DISCUSSION

In this study, we identified a signature of LDN-related genes across various viral and bacterial infections. Our findings indicate that LDNs may play a crucial role in the immune response, especially during acute infections. Our gene expression meta-analysis revealed that genes associated with neutrophil degranulation were significantly upregulated in patients with infections, emphasizing the involvement of LDNs in innate immune processes. Although previous studies have noted the presence of LDNs in severe infections [9, 24], this signature provides a more detailed understanding of LDN biology and its potential impact on immune responses in infectious diseases.

In order to explore the differential regulation of LDN-related genes in acute versus chronic infections and what these patterns suggest about the immune response mechanisms in these conditions, we began our analysis by examining the expression of various genes in 13 PBMC datasets from both acute and chronic infections, including a total of 253 patients and 91 healthy individuals. Our initial analysis of 49 LDN-related genes revealed predominantly increased gene expression in acute infections caused by pathogens such as *S. aureus*, RSV, *E. coli*, *S. pneumoniae*, and Influenza. Conversely, infections like HCV, HIV (chronic), Malaria, and Rotavirus (acute) exhibited mixed expression patterns, while TB and CHB were more likely to show an upregulated profile.

This heterogeneity in gene expression highlights the complexity and adaptive roles that LDNs may play in different infection states [15–21]. These findings probably reflect the multifunctional roles attributed to LDNs in diseases, including their immunosuppressive functions in cancer and infection, mediated by the generation of ROS and release of arginase-1, resulting in inhibition of T-cell proliferation. LDNs may, however, exert functions as pro-inflammatory mediators, promoting NET formation and subsequent endothelial toxicity [15–21].

An important finding was the upregulation of genes such as *LTF*, *OLFM4*, *DEFA4*, *LCN2*, *ELANE*, and *MMP8* in acute infections like *S. aureus*, RSV, *E. coli*, *S. pneumoniae*, and Influenza, indicating a robust antimicrobial response through increased neutrophil degranulation during bacterial and viral infections [25, 30]. This expression may result from the rapid clinical progression and higher severity potential in acute infections, leading to a greater risk of hospitalization. This is linked to a “left shift,” marked by the expansion of immature neutrophils in peripheral blood due to rapid bone marrow mobilization in emergency granulopoiesis [3, 31].

We further explored the LDN-related gene signature and its association with disease severity by analyzing multiple RNA-seq datasets containing information on disease status. This analysis resulted in a list of genes associated with disease severity. Next, we examined a RNA-seq dataset of LDNs classified as the CD16^int^ LDN population, representing immature neutrophils. Comparing these two lists with our previously identified set of 49 LDN genes, we identified at a final list of 17 overlapping genes: *LCN2*, *CAMP*, *MMP8*, *TCN1*, *PGLYRP1*, *ARG1*, *DEFA4*, *CEACAM8*, *PRTN3*, *AZU1*, *LTF*, *HP*, *RNASE3*, *MPO*, *CTSG*, *OLFM4*, and *CEACAM6*. These results provide further evidence that patients with severe acute infections consistently overexpress genes characteristic of immature neutrophils, particularly those in the CD16int LDN population, which possess a band-shaped nucleus.

Several of these 17 genes have been previously associated with infectious conditions, supporting our findings[3, 7, 8, 11, 14, 32, 33]. Moreover, these genes have been linked, either collectively or individually, to the severity of various infectious diseases [14, 29, 32, 34–36]. This association may be connected to the process known as emergency granulopoiesis, or ‘left shift.’

The ‘left shift’ refers to an increase in immature granulocytes, such as band cells, in the peripheral blood [3, 14, 37]. During acute infections, the body initiates emergency myelopoiesis to rapidly produce and release immature neutrophils, including those with a CD16^int^ profile, to meet the heightened demand for immune cells [3, 14, 37]. Clinically, the detection of a ‘left shift’ indicates not only an active immune response but also correlates with the severity and progression of the infection [38], reflecting an active and stressed bone marrow response [3, 30, 37]. Furthermore, the presence of dysfunctional neutrophils in severe infections suggests the activation of multiple deleterious pathways [14, 39]. Understanding the involvement of these 17 genes in the ‘left shift’ process underscores their potential as biomarkers for the host’s immune response during acute infections, highlighting their prognostic significance.

Next, we explored protein abundance in a PBMC proteomics cohort for the 17 identified genes and found 13 LDN-related proteins, with 10 showing significantly higher abundance in sepsis and septic shock patients compared to healthy controls. Our initial hypothesis was that protein abundance in the septic shock cohort would be higher than in sepsis patients, given that septic shock represents a subset of sepsis in which the risk of mortality is substantially increased [40] and that the corresponding genes were found to be upregulated in more severe cases. However, contrary to our expectations, no distinct pattern of LDN-related protein abundance was observed in the septic shock patients.

In contrast, in plasma proteomic studies, some of the proteins listed here have previously been associated with critical illness and mortality in COVID-19 patients [36, 39]. The lack of a clear pattern of association of protein abundance in the most severely ill patients in our proteomics data may be due challenges in correlating transcriptomic data with protein levels [41, 42] or the limited sample size in the PBMC cohort. Therefore, further research with a larger PBMC cohort from septic patients is required to better understand the association between LDN-related proteins and disease severity.

In conclusion, our study identified a common pattern of 17 upregulated LDN genes associated with acute infections and severe clinical outcomes, marking the first time these genes have been collectively presented as a potential signature of LDNs related to disease severity. Although elevated levels of LDN-related proteins were observed in sepsis and septic shock patients, we found no distinct pattern differentiating the two, underscoring the complexities of correlating transcriptomic and proteomic data. Further research with larger cohorts is needed to validate this LDN signature, explore its clinical implications, and understand its translation into protein abundance, potentially paving the way for novel biomarkers to improve prognosis and therapeutic strategies in infectious diseases.

## Acknowledgments and Sources Funding

This work was funded by FAPESP (Grants 2017/21052-0 and 2020/05110-2) to RS. GGFL has a scholarship from FAPESP (2019/20532-3). MAT has a scholarship from FAPESP (2022/12847-7). JVSL has a scholarship from FAPESP (2022/13073-5)

## Authorship Contribution Statement

MAT (Data curation—formal analysis; Investigation; Writing—original draft and Writing— review, editing, and revision)

JVSL (Data curation—formal analysis; Investigation; Writing—original draft and Writing— review, editing, and revision)

GGFL (Conceptualization; Data curation—formal analysis; Investigation; Project Administration/oversight; Writing—original draft and Writing—review, editing, and revision)

RS (Conceptualization; Project Administration/oversight; Writing—original draft and Writing— review, editing, and revision)

## Supplementary Material

### Supplementary Material 1

**Supplementary Table S1.** Additional information about the PBMC microarray datasets selected from GEO

**Supplementary Table S2.** Additional information about the PBMC RNA-seq datasets selected from GEO

**Supplementary Table S3.** MetaVolcanoR vote-counting approach results

### Supplementary Material 2

**Supplementary Table S4**. List of the 49 genes common to both annotations neutrophil degranulation pathway and granule lumen-specific community

**Supplementary Table S5**. Random Effect Model MetaVolcano results for the datasets of patients with acute and chronic infections, filtering for the 49 genes in common between the neutrophil degranulation pathway and granule lumen-specific community

**Supplementary Table S6**. Random Effect Model MetaVolcano results for sensitivity analysis using only microarray data from patients with acute infections

**Supplementary Table S7**. Random Effect Model MetaVolcano results for the analysis involving the 7 RNA-seq datasets with patients stratified by severity

**Supplementary Table S8**. Pairwise comparison of LDN-related protein abundances in sepsis patients and healthy controls

## Data Availability Statement

Data used in this study are freely available at the National Center for Biotechnology Information Gene Expression Omnibus (NCBI GEO) portal: https://www.ncbi.nlm.nih.gov/geo/ and mass spectrometry data were deposited to the ProteomeXchange Consortium via the MassIVE partner repository with the dataset identifier MSV000087733.

## Code availability

All analytic codes are available on reasonable request from the corresponding author.

