## Supplementary Material 2 for "Characterizing a Low-Density Neutrophil gene signature in acute and chronic infections and its impact on disease severity"

**Supplementary Tables**

**Supplementary Table S4**. List of the 49 genes common to both annotations neutrophil degranulation pathway and granule lumen-specific community

| **Gene Symbol** | **Official Full Name** |
| --- | --- |
| *LCN2* | Lipocalin 2 |
| *CAMP* | Cathelicidin antimicrobial peptide |
| *MMP8* | Matrix metallopeptidase 8 |
| *TCN1* | Transcobalamin 1 |
| *PGLYRP1* | Peptidoglycan recognition protein 1 |
| *ARG1* | Arginase 1 |
| *SLPI* | Secretory leukocyte peptidase inhibitor |
| *S100A12* | S100 calcium binding protein A12 |
| *DEFA4* | Defensin alpha 4 |
| *CEACAM8* | CEA cell adhesion molecule 8 |
| *PRTN3* | Proteinase 3 |
| *AZU1* | Azurocidin 1 |
| *LTF* | Lactotransferrin |
| *HP* | Haptoglobin |
| *CD177* | CD177 molecule |
| *RNASE3* | Ribonuclease A family member 3 |
| *TOLLIP* | Toll interacting protein |
| *MPO* | Myeloperoxidase |
| *CTSG* | Cathepsin G |
| *OLFM4* | Olfactomedin 4 |
| *S100A8* | S100 calcium binding protein A8 |
| *SERPINB1* | Serpin family B member 1 |
| *CEACAM6* | CEA cell adhesion molecule 6 |
| *S100A9* | S100 calcium binding protein A9 |
| *RNASE2* | Ribonuclease A family member 2 |
| *ORM1* | Orosomucoid 1 |
| *ITGAM* | Integrin subunit alpha M |
| *ELANE* | Elastase, neutrophil expressed |
| *CTSB* | Cathepsin B |
| *GSDMD* | Gasdermin D |
| *ALOX5* | Arachidonate 5-lipoxygenase |
| *CYBB* | Cytochrome b-245 beta chain |
| *PTAFR* | Platelet activating factor receptor |
| *S100A11* | S100 calcium binding protein A11 |
| *TIMP2* | TIMP metallopeptidase inhibitor 2 |
| *CST3* | Cystatin C |
| *BST1* | Bone marrow stromal cell antigen 1 |
| *CHI3L1* | Chitinase 3 like 1 |
| *GRN* | Granulin precursor |
| *FPR2* | Formyl peptide receptor 2 |
| *PTX3* | Pentraxin 3 |
| *SERPINA1* | Serpin family A member 1 |
| *FPR1* | Formyl peptide receptor 1 |
| *PPBP* | Pro-platelet basic protein |
| *TLR2* | Toll-like receptor 2 |
| *LYZ* | Lysozyme |
| *PLAUR* | Plasminogen activator, urokinase receptor |
| *OLR1* | Oxidized low density lipoprotein receptor 1 |
| *LGALS3* | Galectin 3 |

**Supplementary Table S5**. Random Effect Model MetaVolcano results for the datasets of patients with acute and chronic infections, filtering for the 49 genes common to the neutrophil degranulation pathway and granule lumen-specific community

| **Gene Symbol** | **REM Log2FC** | **REM p-value** |
| --- | --- | --- |
| *TIMP2* | 0.51 | **0.0001** |
| *RNASE2* | 1.22 | **0.0002** |
| *GRN* | 0.51 | **0.0002** |
| *CAMP* | 0.73 | **0.0006** |
| *ELANE* | 0.78 | **0.001** |
| *CEACAM6* | 0.49 | **0.001** |
| *SLPI* | 0.4 | **0.001** |
| *ORM1* | 0.37 | **0.001** |
| *LCN2* | 0.88 | **0.0011** |
| *LTF* | 1.21 | **0.0013** |
| *CEACAM8* | 1.11 | **0.0013** |
| *DEFA4* | 0.91 | **0.0016** |
| *RNASE3* | 0.44 | **0.002** |
| *CTSG* | 0.3 | **0.002** |
| *OLFM4* | 0.96 | **0.0026** |
| *MPO* | 0.58 | **0.003** |
| *ARG1* | 0.48 | **0.003** |
| *AZU1* | 0.46 | **0.003** |
| *MMP8* | 0.73 | **0.0037** |
| *SERPINB1* | 0.33 | **0.004** |
| *PRTN3* | 0.32 | **0.004** |
| *BST1* | 0.67 | **0.01** |
| *HP* | 0.6 | **0.01** |
| *TCN1* | 0.39 | **0.01** |
| *CD177* | 0.3 | **0.01** |
| *TOLLIP* | 0.18 | **0.01** |
| *S100A12* | 0.62 | **0.02** |
| *SERPINA1* | 0.5 | **0.02** |
| *CST3* | 0.44 | **0.02** |
| *S100A9* | 0.5 | **0.04** |
| *PGLYRP1* | 0.26 | **0.04** |
| *LGALS3* | 0.31 | 0.07 |
| *FPR1* | 0.33 | 0.08 |
| *S100A11* | 0.28 | 0.08 |
| *TLR2* | 0.3 | 0.1 |
| *FPR2* | 0.36 | 0.13 |
| *S100A8* | 0.25 | 0.14 |
| *CYBB* | 0.27 | 0.17 |
| *PTX3* | 0.3 | 0.2 |
| *CTSB* | 0.18 | 0.24 |
| *GSDMD* | 0.13 | 0.24 |
| *LYZ* | 0.14 | 0.28 |
| *PLAUR* | 0.4 | 0.29 |
| *ITGAM* | 0.21 | 0.35 |
| *ALOX5* | 0.13 | 0.35 |
| *PTAFR* | 0.11 | 0.36 |
| *CHI3L1* | 0.28 | 0.41 |
| *OLR1* | -0.51 | 0.43 |
| *PPBP* | -0.06 | 0.86 |

**Supplementary Table S6**. Random Effect Model MetaVolcano results for sensitivity analysis using only microarray data from patients with acute infections

| **Gene Symbol** | **Sign consistency** | **REM Log2FC** | **REM p-value** |
| --- | --- | --- | --- |
| *LGALS3* | 8 | 0.730666 | **3.54E-12** |
| *TIMP2* | 8 | 0.692892 | **1.57E-11** |
| *PLAUR* | 8 | 1.217334 | **1.36E-10** |
| *BST1* | 6 | 0.77896 | **6.83E-10** |
| *RNASE2* | 8 | 1.541138 | **5.19E-09** |
| *S100A9* | 6 | 0.718711 | **1.64E-08** |
| *GRN* | 8 | 0.619475 | **5.36E-08** |
| *SERPINB1* | 8 | 0.567798 | **3.88E-07** |
| *PTX3* | 8 | 0.700797 | **6.92E-07** |
| *TLR2* | 6 | 0.71531 | **2.71E-06** |
| *OLR1* | 8 | 0.224422 | **8.00E-06** |
| *SERPINA1* | 8 | 0.702199 | **8.28E-06** |
| *S100A12* | 6 | 0.962518 | **1.96E-05** |
| *FPR1* | 8 | 0.608598 | **2.60E-05** |
| *FPR2* | 8 | 0.776272 | **7.30E-05** |
| *TOLLIP* | 5 | 0.250907 | **7.73E-05** |
| *S100A11* | 6 | 0.559217 | **0.000159** |
| *CEACAM6* | 8 | 0.790276 | **0.001001** |
| *ELANE* | 6 | 1.22127 | **0.001314** |
| *CAMP* | 6 | 1.03542 | **0.002215** |
| *CEACAM8* | 6 | 1.559946 | **0.002232** |
| *LTF* | 6 | 1.614208 | **0.002615** |
| *DEFA4* | 6 | 1.270665 | **0.002694** |
| *LCN2* | 4 | 1.246107 | **0.002703** |
| *PRTN3* | 6 | 0.645292 | **0.002984** |
| *ORM1* | 5 | 0.562037 | **0.003005** |
| *RNASE3* | 6 | 0.632843 | **0.003314** |
| *AZU1* | 6 | 0.800191 | **0.004715** |
| *OLFM4* | 4 | 1.373948 | **0.005097** |
| *CTSG* | 6 | 0.467052 | **0.005842** |
| *MPO* | 6 | 0.845999 | **0.006536** |
| *SLPI* | 6 | 0.541281 | **0.008165** |
| *ARG1* | 6 | 0.818453 | **0.008717** |
| *MMP8* | 6 | 1.126364 | **0.009459** |
| *CST3* | 4 | 0.500928 | **0.01375** |
| *S100A8* | 2 | 0.336559 | **0.015602** |
| *HP* | 6 | 0.75632 | **0.016135** |
| *CD177* | 5 | 0.525696 | **0.019885** |
| *CTSB* | 4 | 0.395236 | **0.021139** |
| *CHI3L1* | 6 | 0.636237 | **0.024102** |
| *CYBB* | 4 | 0.429149 | **0.034985** |
| *PGLYRP1* | 4 | 0.517776 | **0.04299** |
| *ITGAM* | 4 | 0.274703 | 0.063739 |
| *TCN1* | 6 | 0.508905 | 0.076632 |
| *ALOX5* | 2 | 0.130891 | 0.343236 |
| *PTAFR* | 0 | 0.079199 | 0.386008 |
| *LYZ* | 4 | 0.095823 | 0.535675 |
| *PPBP* | 2 | 0.207227 | 0.672506 |
| *GSDMD* | -1 | -0.02264 | 0.790074 |

**Supplementary Table S7**. Random Effect Model MetaVolcano results for the analysis involving the 7 RNA-seq datasets with patients stratified by severity

| **Gene Symbol** | **Sign consistency** | **REM Log2FC** | **REM p-value** |
| --- | --- | --- | --- |
| *LCN2* | 8 | 1.41 | **2.77E-06** |
| *CAMP* | 6 | 1.20 | **5.69E-06** |
| *MMP8* | 5 | 2.23 | **8.91E-04** |
| *TCN1* | 5 | 1.08 | **1.49E-03** |
| *PGLYRP1* | 5 | 1.34 | **2.20E-03** |
| *ARG1* | 5 | 0.75 | **2.75E-03** |
| *SLPI* | 6 | 0.85 | **4.36E-03** |
| *S100A12* | 6 | 1.05 | **4.91E-03** |
| *DEFA4* | 5 | 2.15 | **5.72E-03** |
| *CEACAM8* | 5 | 1.71 | **6.28E-03** |
| *PRTN3* | 5 | 1.87 | **6.35E-03** |
| *AZU1* | 6 | 1.53 | **9.79E-03** |
| *LTF* | 7 | 1.45 | **1.10E-02** |
| *HP* | 6 | 1.10 | **1.20E-02** |
| *CD177* | 3 | 1.65 | **1.86E-02** |
| *RNASE3* | 5 | 0.96 | **2.24E-02** |
| *TOLLIP* | 4 | 0.16 | **2.47E-02** |
| *MPO* | 8 | 0.68 | **2.64E-02** |
| *CTSG* | 3 | 1.99 | **3.07E-02** |
| *OLFM4* | 5 | 1.94 | **3.64E-02** |
| *S100A8* | 4 | 0.71 | **3.72E-02** |
| *SERPINB1* | 6 | 0.40 | **4.08E-02** |
| *CEACAM6* | 3 | 1.49 | **5.01E-02** |
| *S100A9* | 6 | 0.60 | **5.00E-02** |
| *RNASE2* | 4 | 0.31 | 6.41E-02 |
| *ORM1* | 2 | 0.78 | 6.44E-02 |
| *ITGAM* | 4 | 0.43 | 1.09E-01 |
| *ELANE* | 4 | 1.21 | 1.24E-01 |
| *CTSB* | 6 | 0.22 | 1.31E-01 |
| *GSDMD* | 2 | 0.17 | 1.42E-01 |
| *ALOX5* | 6 | 0.38 | 1.54E-01 |
| *CYBB* | 4 | 0.34 | 1.61E-01 |
| *PTAFR* | 4 | 0.35 | 1.80E-01 |
| *S100A11* | 6 | 0.27 | 2.34E-01 |
| *TIMP2* | 6 | 0.23 | 2.41E-01 |
| *CST3* | 0 | -0.19 | 2.44E-01 |
| *BST1* | 4 | 0.26 | 3.03E-01 |
| *CHI3L1* | 3 | 0.27 | 3.13E-01 |
| *GRN* | 4 | 0.15 | 3.29E-01 |
| *FPR2* | 6 | 0.35 | 3.44E-01 |
| *PTX3* | 2 | 0.23 | 4.02E-01 |
| *SERPINA1* | 6 | 0.25 | 4.27E-01 |
| *FPR1* | 4 | 0.24 | 4.83E-01 |
| *PPBP* | 2 | 0.22 | 5.34E-01 |
| *TLR2* | 4 | 0.13 | 5.70E-01 |
| *LYZ* | 2 | 0.10 | 5.75E-01 |
| *PLAUR* | 2 | -0.09 | 6.14E-01 |
| *OLR1* | 1 | -0.26 | 6.78E-01 |
| *LGALS3* | 2 | 0.05 | 6.92E-01 |

**Supplementary Table S8**. Pairwise comparison of LDN-related protein abundances in sepsis patients and healthy controls

| **UniProt ID** | **Gene Symbol** | **Sepsis (n=24)** | **Control (n=9)** | **P-value** |
| --- | --- | --- | --- | --- |
| Q6UX06 | *OLFM4* | 12.91 (2.17) | 9.17 (0.63) | **2.05E-07** |
| P24158 | *PRTN3* | 14.66 (0.79) | 13.13 (0.45) | **1.72E-06** |
| P05164 | *MPO* | 16.22 (0.81) | 14.42 (0.69) | **3.26E-05** |
| P12724 | *RNASE3* | 15.30 (0.87) | 13.71 (0.66) | **6.37E-05** |
| P22894 | *MMP8* | 13.22 (0.91) | 11.87 (0.62) | **1.98E-04** |
| P31997 | *CEACAM8* | 9.12 (1.68) | 6.09 (1.61) | **5.57E-04** |
| P02788 | *LTF* | 15.62 (1.02) | 14.54 (0.58) | **1.52E-03** |
| P80188 | *LCN2* | 10.99 (0.86) | 10.28 (0.39) | **4.94E-03** |
| P20160 | *AZU1* | 10.14 (0.74) | 9.28 (0.65) | **7.27E-03** |
| P49913 | *CAMP* | 12.78 (1.86) | 10.88 (1.85) | **2.56E-02** |
| P00738 | *HP* | 7.11 (1.59) | 6.19 (1.09) | 8.52E-02 |
| P08311 | *CTSG* | 11.75 (0.67) | 11.38 (0.57) | 1.39E-01 |
| P05089 | *ARG1* | 7.40 (1.31) | 6.44 (2.61) | 3.14E-01 |
